# Nanoscale organization of rotavirus replication machineries

**DOI:** 10.1101/445262

**Authors:** Yasel Garcés, José L. Martínez, David T. Hernández, Haydee O. Hernández, Mayra Méndez, Christopher D. Wood, Juan Manuel Rendón Mancha, Daniela Silva-Ayala, Susana López, Adán Guerrero, Carlos F. Arias

## Abstract

Rotavirus genome replication and assembly take place in cytoplasmic electron dense inclusions termed viro-plasms (VPs). Previous conventional optical microscopy studies observing the intracellular distribution of rotavirus proteins and their organization in VPs have lacked molecular-scale spatial resolution, due to inherent spatial resolution constraints. In this work we employed super-resolution microscopy to reveal the nanometric-scale organization of VPs formed during rotavirus infection, and quantitatively describe the structural organization of seven viral proteins and viral dsRNA within and around the VPs. The observed viral components are spatially organized as 6 concentric layers, in which NSP5 localizes at the center of the VPs, surrounded by a layer of NSP2 and NSP4 proteins, followed by an intermediate zone comprised of the VP1, VP2, VP6 proteins and the dsRNA. In the outermost zone, we observed a ring of VP4 and finally a layer of VP7. These findings show that rotavirus VPs are highly organized organelles.

## 1. Introduction

Rotavirus is a non-enveloped virus composed of three concentric layers of proteins that enclose a genome constituted by eleven segments of double stranded RNA (dsRNA) that encode six structural proteins (VP1 to VP4, VP6 and VP7) and six non-structural proteins (NSP1 to NSP6). The inner shell is formed by dimers of VP2 that enclose the viral genome and small amounts of the virus polymerase VP1 and the guanylyltransferase VP3. This nucleoprotein complex constitutes the core of the virus, which is surrounded by an intermediate protein layer of trimers of VP6, to form double-layered particles (DLPs). In the surface of the virion there are two polypeptides, VP7, a glycoprotein that shapes the outer layer of the virus, and VP4, which forms spikes that protrude from the VP7 shell [1]. Replication of the rotavirus genome and assembly of DLPs take place in cytoplasmic electron dense inclusions termed viroplasms (VPs) [1]. Once the double-shelled particles are assembled, they bud from the cytoplasmic VPs into the adjacent endoplasmic reticulum (ER). During this process, which is mediated by the interaction of DLPs with the ER transmembrane viral protein NSP4, the particles acquire a temporary lipid bilayer, modified by VP7, which after being removed in the lumen of the ER by an unknown mechanism, yields the mature triple-layered virions [1]. It has been reported that VP4 is located between the VP and the ER membrane and it is incorporated into triple-layered particles (TLPs) during the budding process and maturation of the virus particle inside the ER [1, 2]. The viral non-structural proteins NSP2 and NSP5 serve a nucleation role that is essential for the biogenesis of VPs [3]. In addition to viral proteins and genomic dsRNA, cellular proteins such as ER chaperones [4], proteins associated with lipid droplets [5], and ribonuclear proteins [6], have been shown to colocalize with VPs.

Several studies have characterized the intracellular distribution of the rotavirus proteins [7, 8, 9, 10]. Immunofluorescence studies, based upon epifluorescence or confocal microscopy, have described the viral proteins that conform the VPs, however the images are inherently diffraction-limited to a spatial resolution in the range of hundreds of nanometers, what precludes identifying the nanoscopic molecular scale organization of VPs [11, 7, 12, 13, 14, 15, 16]. On the other hand, transmission electron microscopy (TEM) studies often provide images with nanometric resolution, nevertheless, immunoelectron microscopy is challenging when looking for the localization of more than a single protein [17, 8, 9]. Over the past 15 years, a variety of super-resolution microscopy (SRM) techniques have been developed to observe subcellular structures beneath the diffraction limit of optical microscopes, with resolutions in the tents of nanometers [18, 19, 20].

In this work, we determined the organization of rotaviral proteins within and around VPs through the “Bayesian Blinking and Blinking” (3B) SRM technique. We developed a segmentation algorithm to automatically analyze and quantify the relative distribution of seven viral proteins and the genomic dsRNA, and propose a model that describes their relative spatial distribution. Also, we present a dependency model that explains the relationship between the viral proteins. This work establishes a clear structural framework for VP organization that future mechanistic and functional studies must take into account, and establishes key methodologies for future investigations on this subject.

## 2. Results

### Qualitative analysis of VP morphology and structure through SRM

Rotavirus VPs are complex signaling hubs composed of viral and cellular proteins, packed together with viral RNAs. By TEM, they roughly resemble circular electrodense structures whose internal components lack an obvious degree of spatial organization [17, 21]. In this work, we determined the relative spatial distribution of VPs components by immunofluorescence and SRM in MA104 cells infected with the rhesus rotavirus strain RRV at 6 hours post-infection (hpi), using protein-specific antibodies. Due to their important role as nucleating factors during VP biogenesis, we selected either NSP2 or NSP5 as spatial relative reference for the distribution of the VP1, VP2 or VP6 proteins. In addition, we used a specific antibody to analyze the distribution of the viral dsRNA genome within the VPs [22]. VPs were optically sectioned through total internal reflection fluorescence microscopy (TIRF), with an excitation depth of field restricted to 200 *nm* from the coverslip. This approach avoids excitation of fluorophores marking structural components located away from this plane, i.e. towards the inner cellular milieu.

Additionally, NSP2 was also co-immunostained with the viral outer layer protein VP4 as well as with the ER resident proteins NSP4 and VP7, all of which have been reported to form separate ring-like structures that closely associate with VPs [7]. In order to gain more insight into the morphogenesis of rotavirus, we analyzed the distribution of both VP7 monomers (VP7-Mon) and trimers (VP7-Tri) since this protein is assembled into virus particles in the latter form [23]. The nanoscale distribution of VPs was then analyzed through 3B-SRM, with improvements in the technique, developed in the present work, to solve nanoscopic structures (“Stochastic model fitted for 3B super resolution microscopy” in Supplementary Information).

By different methods of analysis VPs exhibit roughly a circular shape (Fig. 1A-C). However, unlike the diffraction-limited image (Fig. 1A), in super-resolution microscopy structural details of VP are appreciated, like the different layer distributions of viral components with respect to NSP2 (Fig. 1B and C). In addition to VPs, by diffraction-limited TIRF microscopy we detected in the cytoplasm several small and dispersed puncta of fluorescence (Fig. 1A), and in these images it is also sometimes possible to differentiate the distribution of NSP2 from that of VP4, a closely viroplasm-associated viral protein; see also [7]); in this case, VP4 is detected as a ring-like structure that surrounds the VP. Nevertheless, the small size of the VPs effectively precludes measurement of component distribution for the majority of its structural elements, as their separation is below the spatial resolution of typical optical microscopes. In contrast, images obtained by 3B-SRM do allow the study of the relative distribution of the VP components (Fig. 1B and 1C). In the case of SRM images of VP4 (Fig. 1B), we observed that this protein forms a ring-like structure that does not colocalize with NSP2, and also ribbon-like projections that extend towards the cytoplasm, details that were not apparent in images captured with conventional fluorescence microscopy (Fig. 1A). Additionally, we observed that the small puncta of proteins detected in the cytoplasm were in fact ribbon-like structures composed of various viral proteins that may represent different organization forms of the viroplasmic proteins (Fig. 1B). In this regard, it is interesting to note that both NSP2 and VP4 have been reported to have at least two different intracellular distributions [7, 24, 14].

**Figure 1.**
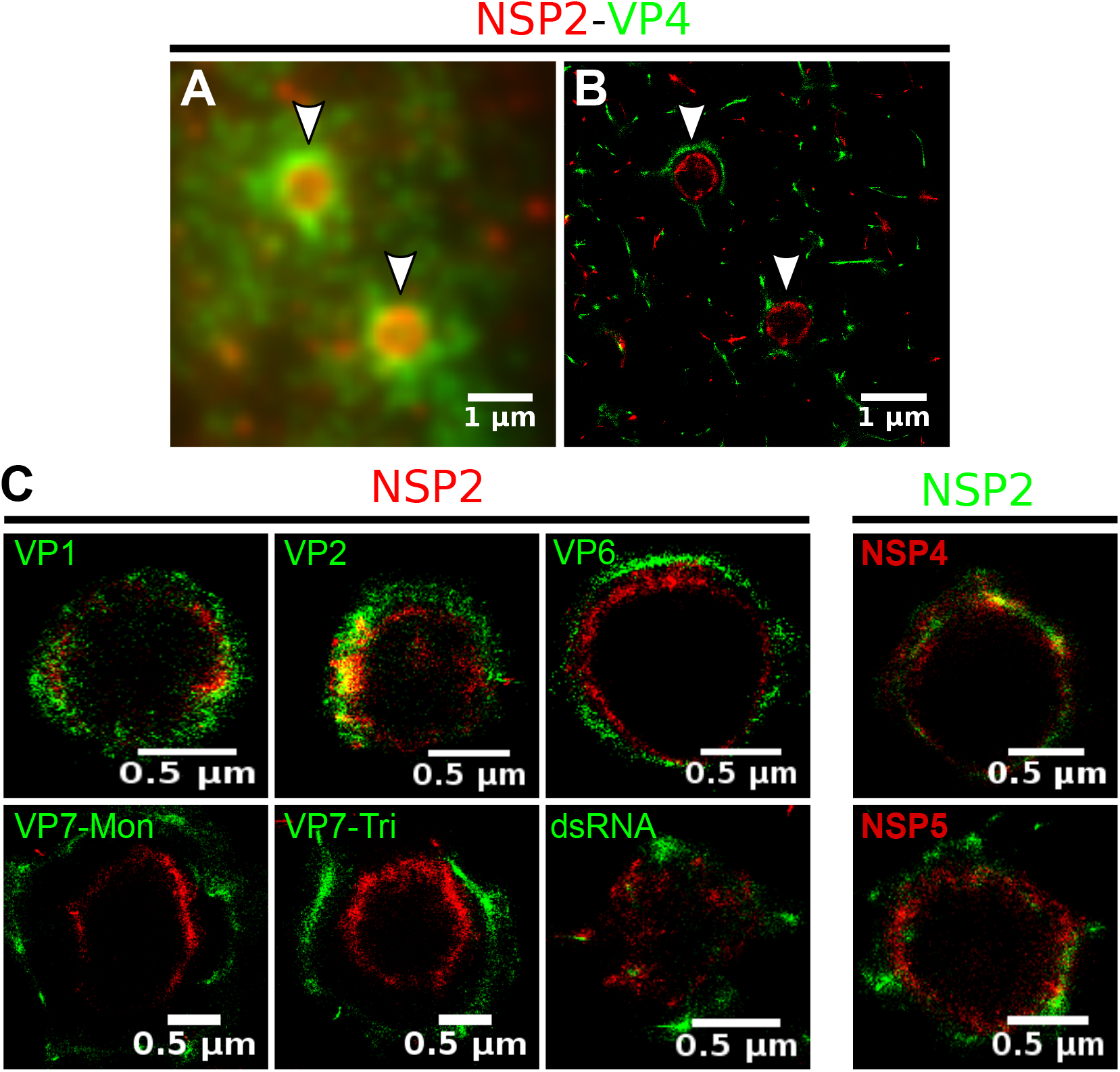
Relative distribution of viral components in rotavirus-VPs. RRV-infected MA104 cells (6hpi) were fixed and processed for this section through immunofluorescence microscopy. **A)** Diffraction-limited image of VPs (white arrows). **B)** 3B-SRM image reconstructed from A. **C)** 3B-SRM images of individual VPs labeled with different antibodies (see Methods).

An examination of 3B-SRM images of VPs (Fig. 1B and 1C) revealed that the viral components form ring like structures within the VPs and are arrayed as rather discrete concentric layers. As seen in Fig. 1C, we find that although the structural proteins VP1, VP2 and VP6 partially overlap in position with NSP2, the bulk of the proteins form separate and distinct layers. Also, the monomeric as well as the trimeric forms of VP7 are clearly distinguished from NSP2, forming an outer ring. Of interest, the spatial distribution of NSP4 colocalized with that of NSP2, an unexpected result since, as mentioned, NSP4 is an ER integral membrane protein (see the Discussion section), and as such it was expected to colocalize with VP7 rather than with an internal viroplasmic protein [9]. With regard to NSP5, it was observed distributed inside the ring formed by NSP2 (Fig. 1C). Finally, the dsRNA was observed forming small clusters at the periphery of NSP2, although these clusters did not form a continuous ring-like structure as for the rest of the viral components; they were, however, concentrated at an intermediate distance from NSP2 and VP4. It is likely that the signal detected with the antibody to dsRNA represents replication-intermediate complexes where the dsRNA might be accessible to interaction with the antibody.

### Quantitative characterization of VPs structure by a novel segmentation algorithm

A qualitative analysis of the distribution of the VP components through 3B-SRM suggested that these are arranged as concentric spherical shells; thus, we set out to quantitatively validate the circularity of the VP shape. For this, we developed a segmentation algorithm based on a least squares approach, which we called “Viroplasm Direct Least Squares Fitting Circumference” (VP-DLSFC) (see “Segmentation Algorithm” in Supplementary Information), to measure the spatial distribution of the components within individual VPs by adjusting concentric circumferences. This method is automatic, deterministic, easy to implement, and has a linear computational complexity. The performance of VP-DLSFC was tested on approximately 40,000 “ground truth” (GT) synthetic images, showing a high robustness to noise and partial occlusion scenarios. Additionally, we compared our method with two other alternative methods [25], and our approach displayed an improved performance (data not shown, see “Algorithm Validation” in Supplementary Information).

Based on this new algorithm, we find that the mean radius of the NSP5 distribution was smaller than that of NSP2, suggesting that NSP5 is located in the innermost section, as a component of the core of VPs (Fig. 2A). On the other hand, the distribution of the structural proteins VP1, VP2 and VP6 exhibit slightly larger mean radii than that of NSP2, and are thus primarily localized in a zone surrounding NSP2. Continuing further towards the outer regions of the VP, we observed a region occupied by the dsRNA and the spike protein VP4. Finally, different forms of VP7 (VP7-Mon and VP7-Tri) were located together, close to the most external region of the VPs (Fig. 2A). The distribution of the glycoprotein NSP4 showed a similar mean radius to that of NSP2 (around 0.4*μm*) suggesting, as described above, that these two proteins are located in the same structural layer of the VP (Fig. 2A).

**Figure 2.**
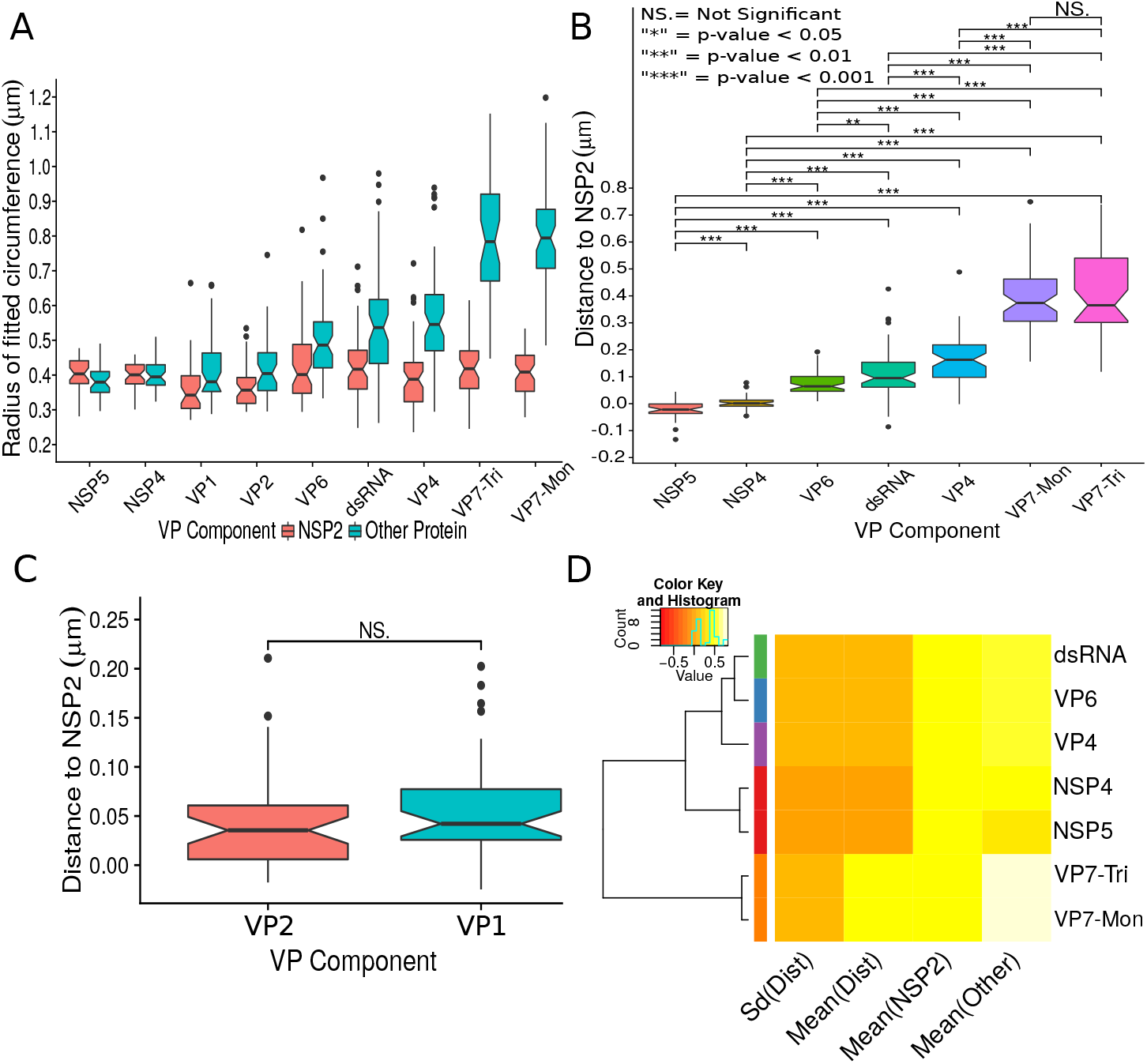
Exploratory analysis of the results obtained by the algorithm VPs-DLSFC. **A)** Boxplot for the radius of the fitting circumferences. In each experimental condition we plot two boxes, the red box is for the radius of NSP2 (reference protein), and the blue box represents the radius of the accompanying VP components (names in x-axis). **B)** Boxplot and results of the Mann-Whitney hypothesis test for the distance between each viral element and NSP2. Each combination of the Mann-Whitney test is linked by a line, and the result of the test it is above the line. Note that this test reports significant differences between the distribution of the distance to NSP2 of two different VP components. **C)** Distance of VP1 and VP2 to NSP2 and result of the Mann-Whitney test. Because the distributions of NSP2 in combination with VP1 and VP2 are statistically different to the other NSP2 distributions (see Figure S7B), we show these two cases independently in this exploratory analysis. **D)** Hierarchically clustered heatmap for the standard deviation of the distance to NSP2, the mean distance to NSP2, the mean radius of NSP2, and the mean radius of the accompanying protein layers, NSP5, NSP4, VP6, VP4, VP7 and the dsRNA.

In order to confirm our preliminary observations and clarify the nanoscopic organization of VPs, we evaluated the relative separation between NSP2 and each accompanying protein. Again, the results show a remarkable degree of organization in the structure of the VP (Fig. 2B). As predicted from Fig. 2A, we found that NSP5 is located in the internal part of the VP, in close proximity (≈ 0.05*μm*) to the area occupied by proteins NSP2 and NSP4, which themselves show the closest association. After the NSP2-NSP4 region, VP6 occupies a middle region at ≈ 0.05*μm* from NSP2, followed by the dsRNA and the VP4 protein, which were located at a distance of ≈ 0.1*μm* and ≈ 0.18*μm*, respectively. Finally, the VP7-Mon and VP7-Tri were situated at ≈ 0.38*μm* from NSP2 (Fig. 2B). A Mann-Whitney test showed that the distances of the various viral components in relation to NSP2 were significantly different (Fig. 2B), suggesting that they are situated in specific areas of the VPs. The two forms of VP7 were located at the same distance to NSP2, suggesting that the formation of trimers of VP7 takes place at the ER membrane, where the VP7 monomers should also be located. Note that in Fig. 2B the relative distance of VP1 and VP2 to NSP2 was not included, since the radii obtained for NSP2 in these two combinations were significantly smaller than those found when it was determined in combination with the other VP components (see “Supplementary Exploratory Analysis”). In addition to this, we found no significant differences between the distance of both VP1 and VP2 to NSP2 (Fig. 2C). Nonetheless, based on the inferential analysis, we could place these two proteins in the same layer as VP6 (see below).

Next, through a hierarchical cluster analysis, we studied the relationship between the components of the VP, taking into account multiple variables at the same time, like the mean distance to NSP2 [“Mean(Dist)”], the standard deviation of the distance to NSP2 [“Std(Dist)”], the mean radius of NSP2 [“Mean(NSP2)”], and the radii of the other proteins [“Mean(Other)”] (Fig. 2D). As we are considering the distance to NSP2, VP1 and VP2 were not included in this analysis. The first level of the hierarchical clustering (Fig. 2D, left) partitioned the VPs and the surrounding proteins in 5 clusters, composed by {NSP4, NSP5}, VP6, dsRNA, VP4 and {VP7-Mon, VP7-Tri}. The next levels indicate that VP6 and the dsRNA are clustered in a second group, and VP4 was merged to {VP6, dsRNA} in the third level. The subsequent groups in the clustering analysis indicate that VP7 remains as an independent layer with respect to the other proteins.

### The relative spatial organization of VPs is maintained regardless their size

The scatterplot between the radius of the spatial distribution of NSP2 (independent variable, x-axis) and the radius of the distribution of other viral components (response variable, y-axis) showed a strong linear relationship (Fig. 3A). The distribution of NSP5 grows 0.87*μm* for each 1*μm* increase in the radius of NSP2 (slope interpretation), whereas the radius of the distribution of NSP4 increases 0.99*μm* (Fig. 3B). These findings indicate that NSP5 is distributed in a proportionally smaller region than NSP2 regardless of the absolute size of the VP, supporting our observation that NSP5 is a constituent of the core of the VP. Moreover, the fact that the increase in the radius of the fitted distribution of NSP4 is directly proportional to the same parameter measured for NSP2 supports the idea that these proteins are both constituents of a putative second layer. VP1, VP2 and VP6 exhibit similar slopes which diverge between 0.03 and 0.05*μm* (Fig. 3B and Table S6); thus, these results confirm that VP1, VP2 and VP6 are components of the same layer in the VPs which, from the data in Fig. 3, is located just after the layer of NSP2 and NSP4. Finally, as noted in our quantitative analysis, dsRNA, VP4 and VP7 form consecutive external layers with a slope of 1.25, 1.39 and 1.94*μm*, respectively (Fig. 3B and Table S6). These findings indicate that the spatial distribution of the viral components in the VPs and in the surrounding areas is conserved regardless of their absolute size, and also form the basis of a predictive model, where, for a given radius of distribution of NSP2, it is possible to predict the radii of the remaining VP components (NSP5, NSP4, VP1, VP2, VP6, dsRNA) and of VP4 and VP7 proteins. This predictive model is available as a web app here^1^. The mathematical details and the residual analysis that validate these linear models are available in the Supplementary Information, section “Linear dependency between the viral components”, Table S6 and Figure S8.

**Figure 3.**
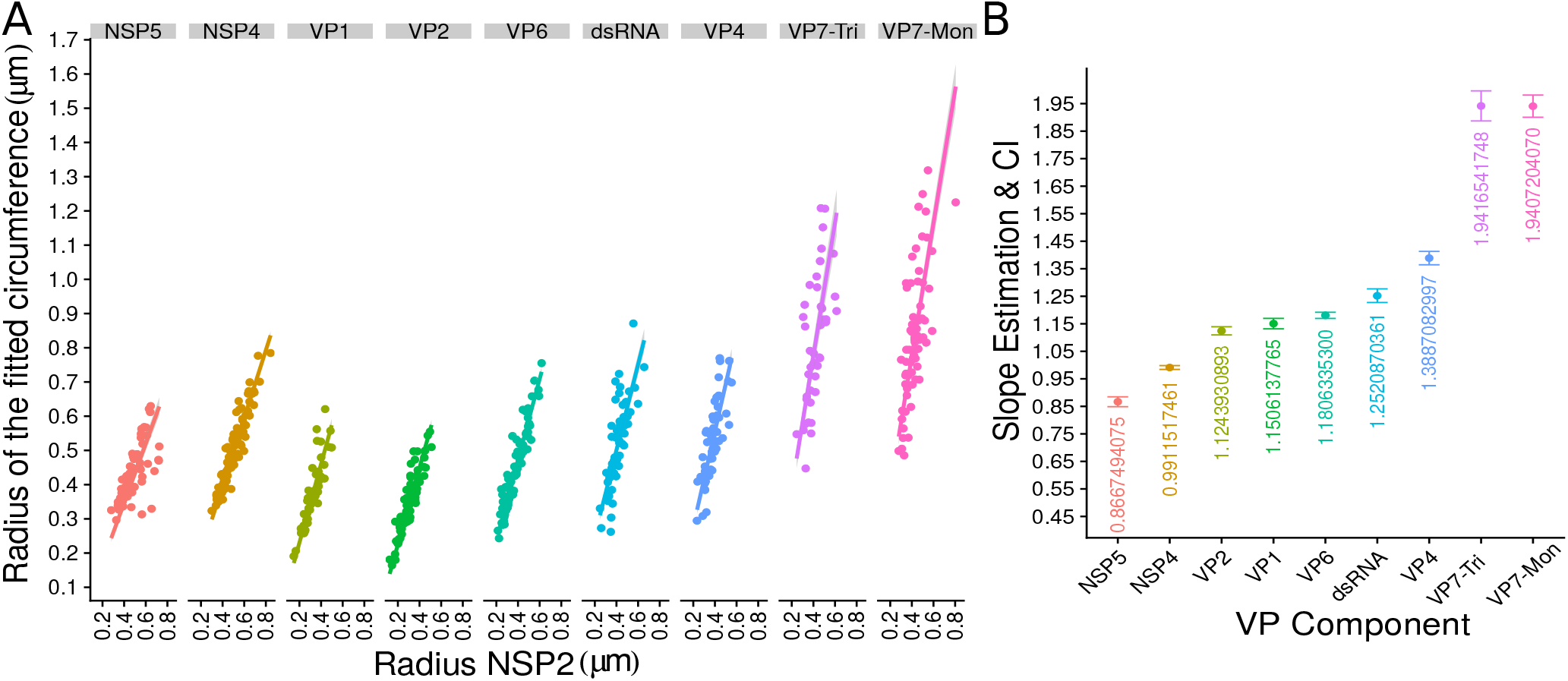
The organization of VPs scales with its size. **A)** Simple linear regression analyses for each component combination (9 subpanels). In all subpanels, the x-axis represents the radius of the distributions of NSP2, and the y-axis the radius of the distribution of the accompanying VP component. The 95% confidence interval, marked in grey, is imperceptible due to goodness of fit of the linear regression (solid line). **B)** Slope and confidence interval for each linear regression model (dependent variables in x-axis). The slopes values were shown under each confidence interval for a better interpretation of the graph.

### The structural organization of VPs is independent of the reference protein chosen for pairwise comparison

In order to confirm the observed structural organization of VPs, we analyzed two more experimental conditions in which we chose a different reference protein for pairwise comparisons. The first was based on the distribution of NSP5 and its comparison with the relative localizations of VP6 and VP4, and the second considered NSP4 as the reference protein to compare with the distribution of VP6. We found that both analyses produced an identical structural organization for the VPs, with a comparative localization error of approximately 0.05*μm* between models (close to the effective resolution limit of the 3B algorithm; see “NSP5 and NSP4 as reference proteins” in Supplementary Information). An extensive quantitative validation regarding the congruence between the NSP2, NSP5 and NSP4 models is available in the Supplementary Information.

Based on our extensive quantitative, descriptive and inferential statistical analyses, we propose that the VP and the surrounding viral proteins form an ordered biological structure composed of at least 6 concentric layers organized as depicted in Fig. 4. In this structure, NSP5 constitutes the innermost layer, followed by a {NSP2-NSP4} layer. Then, there is a layer composed by {VP1-VP2-VP6} and three consecutive external layers formed by a discontinuous layer of dsRNA, and two layers formed by VP4 and VP7.

**Figure 4.**
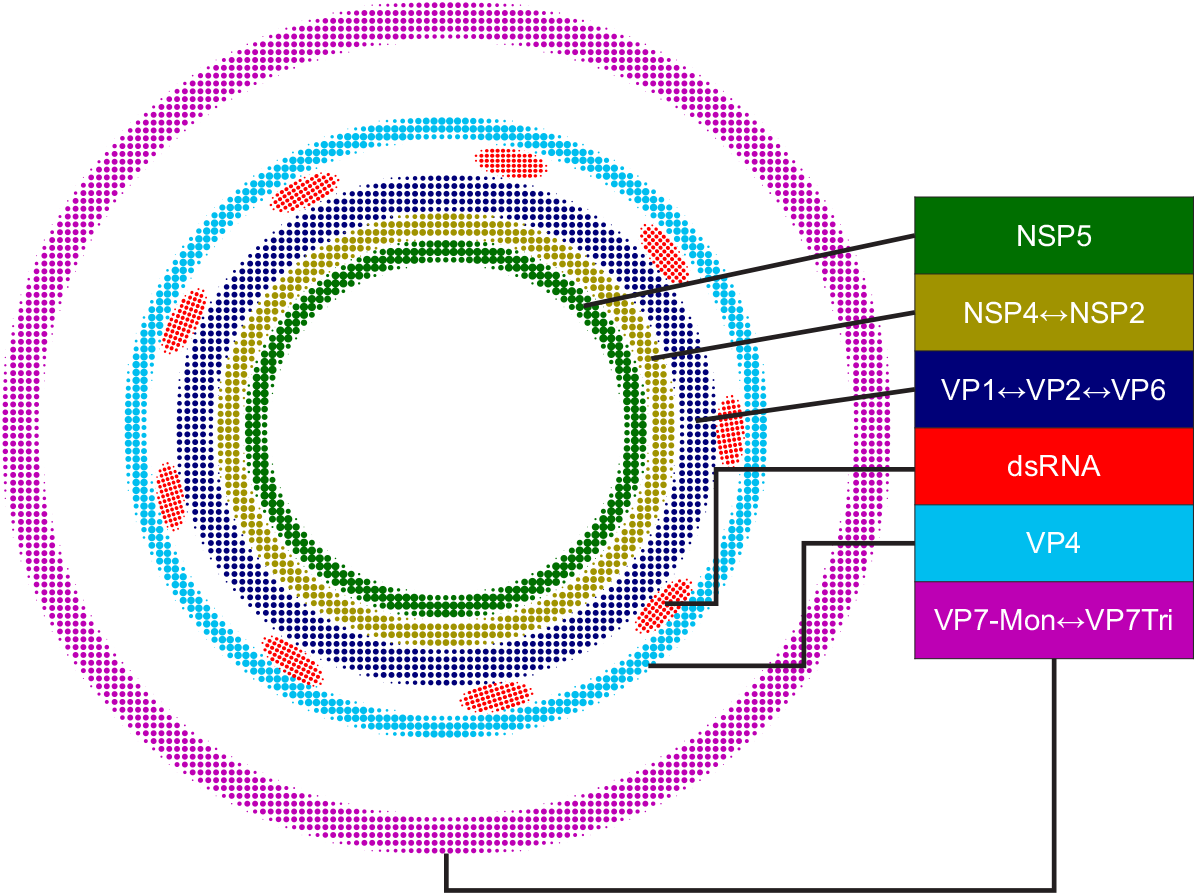
Relative structural distribution of VP components. The radii of the circumferences maintain the relative values determined for the different VP layers.

## Discussion

VPs have been previously studied using electron and fluorescence microscopy, however, due to the limited resolution of classic fluorescence microscopy techniques, and the difficulty of analysis of immunoelectron microscopy, the existence of any complex structural organization of the viral elements inside the VPs has not been reported. In recent years, the development of SRM has facilitated research into the nanoscale organization of a diverse range of cellular structures [26, 27], however, until now SRM had not been applied to study the replication cycle of rotavirus. In this work, for the first time, we visualized and determined quantitatively the location of several protein and nucleic acid components of VPs, what led us to propose a detailed model of the VP that should be of great value for understanding virus morphogenesis.

The quantification of the viral protein distribution within the VPs was possible thanks to a novel segmentation algorithm (VPs-DLSFC) that was proven to be robust and efficient in noisy and partial occlusion scenarios. The manual pre-segmentation step of this algorithm was necessary in our case, because we did not want to introduce any bias in the isolation of the VPs through an automatic approach. Setting aside the manual pre-segmentation step, the VPs-DLSFC algorithm is automatic, deterministic, non-iterative and has a linear computational complexity.

Previous reports have suggested that VPs have a spherical-like structure [12, 28, 29]; in this study we confirmed this suggestion by comparing the VPs-DLSFC approach with a similar approach based on an ellipse adjustment [30]. The results show no significant statistical differences between these two models, and as consequence we can confidently model the structure of the VPs as a circumference. We also ratified that the center displacement of the circumferences that adjust two paired proteins are not statistically different. This approach is simple, computationally efficient and easy to interpret, but also could be a great departure point to more specific investigations, for example, as initial solution for the quantification of topological changes in the shape of the VPs through level-set [31] or region growing methods [32, 33].

Our study indicates that the viral components in the VPs, as well as VP4 and VP7, are arranged as largely discrete concentric layers. This organization, however, does not preclude the interaction between VP components that in the proposed model have been located in separate layers since, for instance, NSP5 has been shown by different biochemical methods to interact with NSP2 [12, 34, 35], VP1 [35], VP2 [36], and dsRNA [37].

Previous studies based on conventional microscopical techniques have reported that NSP5 and NSP2 colocalize [7, 12, 3]; in contrast, we found that although NSP5 and NSP2 are located in close proximity, their positions in the VP were separable. This difference is attributable to the increased spatial resolution in the final image created by the super-resolution techniques employed in our study. Here, NSP5 was found to represent the innermost layer of the VPs, suggesting that this protein might serve as the core scaffold upon which the subsequent viroplasmic proteins are assembled to form the VPs. This finding contrasts with a report by Eichwald et al. [12] that described that NSP5 locates to a region external to NSP2. In addition to the superior spatial resolution obtainable through 3B-SRM, compared to the traditional confocal microscopy employed in the previous report, the difference might be due to the fact that in our study we characterized the endogenous structures produced during virus replication, while Eichwald et al. characterized VP-like structures formed by transiently expressed proteins fused to GFP.

Immediately outside the NSP5 core, we observed a layer composed of NSP2 and NSP4 proteins. The finding that NSP4 is located in the inner part of VPs was unexpected, since it is known that NSP4 is an integral membrane protein of the ER and it has been reported that functions as a receptor for the new DLPs located at the periphery of the VPs, during their budding towards the lumen of the ER [38, 8, 9, 39]. Furthermore, it has been shown that NSP4 associates with VP4 and VP7 to form a hetero-oligomeric complex that could be involved in the last steps of rotavirus morphogenesis [40]. Based on these findings, NSP4 was expected to locate close to VP4 and VP7, in the surroundings of the VP. On the other hand, and in line with our observations, previous confocal microscopy studies have shown that a portion of NSP4 also shows a limited colocalization with NSP2 [7].

The dual location of NSP4 as an integral glycoprotein of the ER membrane and as internal to VPs, as our results indicate, is not easy to reconcile; however, these observations could be explained either if part of the ER membrane is embedded inside the VP, or if one of the described cytosolic, soluble forms of NSP4 is an integral component of VPs [41, 42]. The location of NSP4 internal to VPs could also explain the observation that knocking-down the expression of NSP4 by RNA interference significantly reduces the number and size of VPs present in the cell, as well as the production of DLPs [43]. That study also showed that during RNAi inhibition of NSP4 expression the NSP2 and NSP5 proteins maintained an intracellular distribution restricted to VPs, while the VP2, VP4, VP6 and VP7 proteins failed to locate to VPs. Based on these observations, it is tempting to suggest that, in addition to the role NSP4 has on the budding of DLPs into the ER lumen, it may also play an important role as a regulator of VP assembly.

After the NSP2/NSP4 layer, we observed a middle zone composed of the structural proteins VP1, VP2 and VP6. Their location in the same zone is expected given their close association in the assembled DLPs [1]. Also, the fact that VP1, VP2 and VP6 form a complex with NSP2 that has replicase activity [44], suggests that the production of new DLPs could take place in this zone of the VP. This observation is in agreement with the location of the dsRNA layer positioned next to VP1, VP2 and VP6.

Finally, we found that VP4 and VP7 conform independent layers just external to the viroplasmic proteins. The position of these two proteins agrees with the proposed model of rotavirus morphogenesis in which VP4 is assembled first on DLPs, and subsequently VP7 binds the particles and locks VP4 in place [45]. Furthermore, the fact that VP7-Mon and VP7-Tri occupied the same layer in our model indicates that in the ER sites into which the DLPs bud, VP7 is already organized as trimers, which are subsequently assembled into the virus particles. Of interest, VP4 has been reported to exist in two different forms in infected cells. One of them is associated with microtubules [24], while the other one has been reported to be found between the VP and the ER membrane [7]. In this regard, based on our findings, we suggest that the latter form of VP4 can be actually considered as an integral component of the VP. Several studies have found the presence of different cellular proteins and lipids in association to VPs [4, 5, 6], it will be interesting to study the relative localization of this components using the methodologies described here.

## 3. Methods and Materials

### Cell and virus

The rhesus monkey kidney epithelial cell line MA104 (ATCC) was grown in Dulbecco’s Modified Eagle Medium-Reduced Serum (DMEM-RS) (Thermo-Scientific HyClone, Logan, UT) supplemented with 5% heat-inactivated fetal bovine serum (FBS) (Biowest, Kansas City, MO) at 37*°*C in a 5% CO_2_ atmosphere. Rhesus rotavirus (RRV) was obtained from H. B. Greenberg (Stanford University, Stanford, Calif.) and propagated in MA104 cells as described previously [46]. Prior to infection, RRV was activated with trypsin (10 *μg/ml*; Gibco, Life Technologies, Carlsbad, CA) for 30 min at 37*°*C.

### Antibodies

Monoclonal antibodies (MAbs) to VP1, VP2(3A8), VP4 (2G4), VP6 (255/60), VP7 (60) and VP7 (159) were kindly provided by H. B. Greenberg (Stanford University, Stanford, CA). The rabbit polyclonal sera to NSP2, NSP4 and NSP5, and the mouse polyclonal serum to NSP2 were produced in our laboratory. Mouse monoclonal antibody to dsRNA (SCICONS J2) was purchased from English and Scientific Consulting Kft, Hungary (SCICONS). Goat anti-mouse Alexa-488- and Goat anti-rabbit Alexa-568-conjugated secondary antibodies were purchased from Molecular Probes (Eugene, Oreg.).

### Immunofluorescence

MA104 cells grown on glass coverslips were infected with rotavirus RRV at a multiplicity of infection (MOI) of 1. Six hours post infection, the cells were fixed with and processed for immunofluorescence as described [47]. Finally, the coverslips were mounted onto the center of glass slides with storm solution (1.5% glucose oxidase +100 mM *β* -mercaptoethanol) to induce the blinking of the fluorophores [48, 49].

### Set up of the Optical Microscope

All super-resolution imaging measurements were performed on an Olympus IX-81 inverted microscope configured for total internal reflection fluorescence (TIRF) excitation (Olympus, cellTIRFTM Illuminator). The critical angle was set up such that the evanescence field had a penetration depth of ~200 nm (Xcellence software v1.2, Olympus soft imaging solution GMBH). The samples were continuously illuminated using excitation sources depending on the fluorophore in use. Alexa Fluor 488 and Alexa Fluor 568 dyes were excited with a 488 nm or 568 nm diode-pumped solid-state laser, respectively. Beam selection and modulation of laser intensities were controlled via Xcellence software v.1.2. A full multiband laser cube set was used to discriminate the selected light sources (LF 405/488/561/635 A-OMF, Bright Line; Semrock). Fluorescence was collected using an Olympus UApo N 100*x*/1.49 numerical aperture, oil-immersion objective lens, with an extra 1.6x intermediate magnification lens. All movies were recorded onto a 128×128-pixel region of an electron-multiplying charge coupled device (EMCCD) camera (iXon 897, Model No: DU-897E-CS0-#BV; Andor) at 100 nm per pixel, and within a 50 ms interval (300 images per fluorescent excitation).

### Bayesian Analysis of the Blinking and Bleaching

Sub-diffraction images were derived from the Bayesian analysis of the stochastic Blinking and Bleaching of Alexa Fluor 488 dye [20]. For each super-resolution reconstruction, 300 images were acquired at 20 frames per second with an exposure time of 50 ms at full laser power, spreading the bleaching of the sample over the length of the entire acquisition time. The maximum laser power coming out of the optical fiber measured at the back focal plane of the objective lens, for the 488 nm laser line, was 23.1 mW. The image sequences were analyzed with the 3B algorithm considering a pixel size of 100 nm and a full width half maximum of the point spread function of 270 nm (for Alexa Fluor 488), measured experimentally with 0.17*μm* fluorescent beads (PS-SpeckTM Microscope Point Source Kit, Molecular Probes, Inc.). All other parameters were set up using default values. The 3B analysis was run over 200 iterations, as recommended by the authors in [20], and the final super-resolution reconstruction was created at a pixel size of 10 nm with the ImageJ plugin for 3B analysis [50], using parallel computing as described in [51]. The resolution increase observed in our imaging set up by 3B analysis was up to 5 times below the Abbe’s limit (~50 nm). The resolution provided by 3B was improved by computing the photo-physical properties of Alexa Fluor 488, and Alexa Fluor 568 dyes, which were provided to 3B algorithm, as an input parameter which encompass the probability transition matrix between fluorophore’s states. The method was validated with 40 nm gattapaint nanorules (PAINT 40RG, gattaquant,Inc.) labeled with ATTO 655 / ATTO 542 dyes (see “3B Algorithm” in Supplementary Information).

### Code and Statistical Analysis

The segmentation algorithm (VPs-DLSFC) was developed in Matlab R2018a (9.4.0.813654) software. A detailed explanation of each the developed methods is available in Supplementary Information. Statistical analysis were performed using R version 3.4.4 (2018-03-15) software. All the codes are available in (public link after the work’s acceptance) only for academic research purposes.

## 4. Acknowledgments

Y.G. received a postdoctoral fellowship from DGAPA-UNAM at the Institute of Biotechnology (IBt-UNAM). A.G. thanks DGTIC-UNAM for generous computing time on the Miztli supercomputer (Grant numbers: SC15-1-IR-89; SC16-1-IR-102). J.L.M. and D.T.H. are recipients of scholarships from CONACyT. H.O.H. received a grant from the Programa de Apoyo a Proyectos de Investigación e Innovación Tecnológica (PAPIIT-UNAM), IN202312. Microscopy equipment was provided and maintained through CONACYT grants 123007, 232708, 260541, 280487, 293624. A.G. thanks CONACyT (No. 252213) and DGAPA-PAPIIT (No. IA202417), S.L. and C.F.A. thank DGAPA-PAPIIT grant IG200317 for funding. We are thankful to IBt-UNAM for providing access to the computer cluster and to Jerome Verleyen for his support while using it. We are also thankful to Arturo Pimentel, Andrés Saralegui and Xochitl Alvarado from LNMA-UNAM for their helpful discussions. The funders had no role in study design, data collection and interpretation, or the decision to submit the work for publication.

## Footnotes

1 This is a primary version of the app; since the most recent version includes information related to this work, we will deliver the full app after acceptance of this paper.

